# Response variability and population coupling of cortical synaptic inputs are strongly influenced by network properties

**DOI:** 10.1101/087031

**Authors:** Nathaniel C. Wright, Mahmood Hoseini, Tansel Baran Yasar, Ralf Wessel

## Abstract

The highly variable spiking of a cortical neuron is “coupled” to that of other neurons in the network. This has implications for sensory coding, and appears to represent a fundamental property of cortical sensory processing. To date, most studies of population coupling have focused on recorded spiking activity, an approach that suffers from several confounding issues. Moreover, the contributions of various network properties to population coupling are largely unexplored. To this end, we recorded the membrane potential (V) and the nearby LFP in the visual cortex of the turtle *ex vivo* wholebrain preparation during ongoing and visually-evoked activity. We used an algorithm to infer the excitatory conductance (g) from V, and calculated the g-LFP coupling. We found that g-LFP coupling was highly variable across neurons, and increased following visual stimulation before relaxing to intermediate values. To investigate the role of the network, we implemented a driven small-world network of leaky integrate-and-fire neurons. This model reproduces the large across-trial response variability and g-LFP coupling dynamic, and suggests crucial roles for anatomical and emergent network properties.

---

Cortical neuron sensory responses are remarkably variable across trials(Britten et al. 1993; Carandini 2004; Scholvinck et al. 2015). Because this variability tends to be correlated between pairs of nearby neurons (see Kohen, Cohn 2011 and Doiron, et al. 2016 for reviews), it likely influences population coding of sensory information(Abbott and Dayan 1999; Averbeck, Latham, and Pouget 2006; Shadlen and Newsome 1998). With advances in recording techniques, it has become increasingly obvious that single-neuron variability reflects fluctuations that are shared across large regions of cortex(Lin et al. 2015; Okun et al. 2015; Scholvinck et al. 2015). That is, sensory input interacts with intrinsic cortical activity, with global cortical fluctuations influencing single-neuron responses. Appropriately, a recent study has introduced the term “population coupling” to describe this relationship(Okun et al. 2015). This and other studies have shown that the coupling of spiking activity is remarkably diverse across neurons (likely reflecting connectivity(Okun et al. 2015; Pernice et al. 2011)), yet can also change with sensory stimulation(Haider, Schulz, and Carandini 2016; Tan et al. 2014) and network state(Haider, Schulz, and Carandini 2016; Okun et al. 2015; Scholvinck et al. 2015). Moreover, the effects of this globally-derived input (i.e., additive vs. multiplicative response gain) may reflect specific mechanisms by which feedback exerts its influence on the response(Larkum 2013; Reynolds and Heeger 2009). This rich dependence of single-neuron responses on anatomical and emergent network properties appears to represent a fundamental principle of cortical function, and is only beginning to be explored. Here, we investigate three questions vital to a better understanding of cortical variability and its effects on sensory coding. (1) What is the nature of response variability in cortical microcircuits? (2) How strongly are single-neuron synaptic input fluctuations coupled with those of the local population? (3) To what degree are the dynamics of response variability and population coupling determined by the cortical network, and what are the relevant network parameters?

While spike-based studies have yielded many important insights, this approach has two inherent shortcomings. First, it excludes the vast majority neurons, which are sparse-spiking(Henze et al. 2015; O’Connor et al. 2010; Shoham, O’Connor, and Segev 2006; Thompson and Best 1989) and therefore yield unreliable statistics for the analysis of correlated variability(Cohen and Kohn 2011) (**Figure 1a, b**). Second, it involves sampling populations of neurons that are visible to the experimentalist, but which may not represent relevant or complete cortical microcircuits. Patch clamp recordings of synaptic inputs represent one solution to these two problems(Shoham, O’Connor, and Segev 2006). First, when the recorded neuron is viewed as a component member of the network, the subthreshold inputs provide a measure of activity that is agnostic to output spike rate. A second perspective, motivated by anatomical connectivity, recognizes the neuron as a “device” that samples an enormous and extremely relevant pool of presynaptic neurons. Thus, subthreshold recordings allow the experimenter to “tap into” the cortical circuitry itself, and infer response properties (e.g., variability) of these large populations(Ikegaya et al. 2004; MacLean et al. 2005; Mokeichev et al. 2007) (**Figure 1a, b**). Despite the potential of this technique, it is rarely implemented *in vivo*; it is difficult to obtain stable patch clamp recordings of cortical sensory responses, and spatially-extended cortical pyramidal neurons confound the interpretation of voltage clamp data(Armstrong and Gilly 1992; Koch 2004; Spruston et al. 1993). Here, we overcome these challenges to address the first two questions above. First, we recorded subthreshold membrane potential visual responses from cortical pyramidal neurons in the turtle eye-attached wholebrain *ex vivo* preparation (**Figure 1c**). We then applied a recently-developed algorithm(Yaşar, Wright, and Wessel 2016) to infer the excitatory synaptic conductance (g) from V (**Figure 1c**), and analyzed the response variability in g. Finally, we calculated the correlated variability for g and the nearby local field potential (LFP). We found that visual stimulation evoked significant increases in g and LFP variability. The variability was typically large relative to the average response, and was a mix of multiplicative and additive noise. Across the population of cells, g-LFP correlated variability (CC) was highly variable, and transiently increased with visual stimulation.

**Figure 1.**
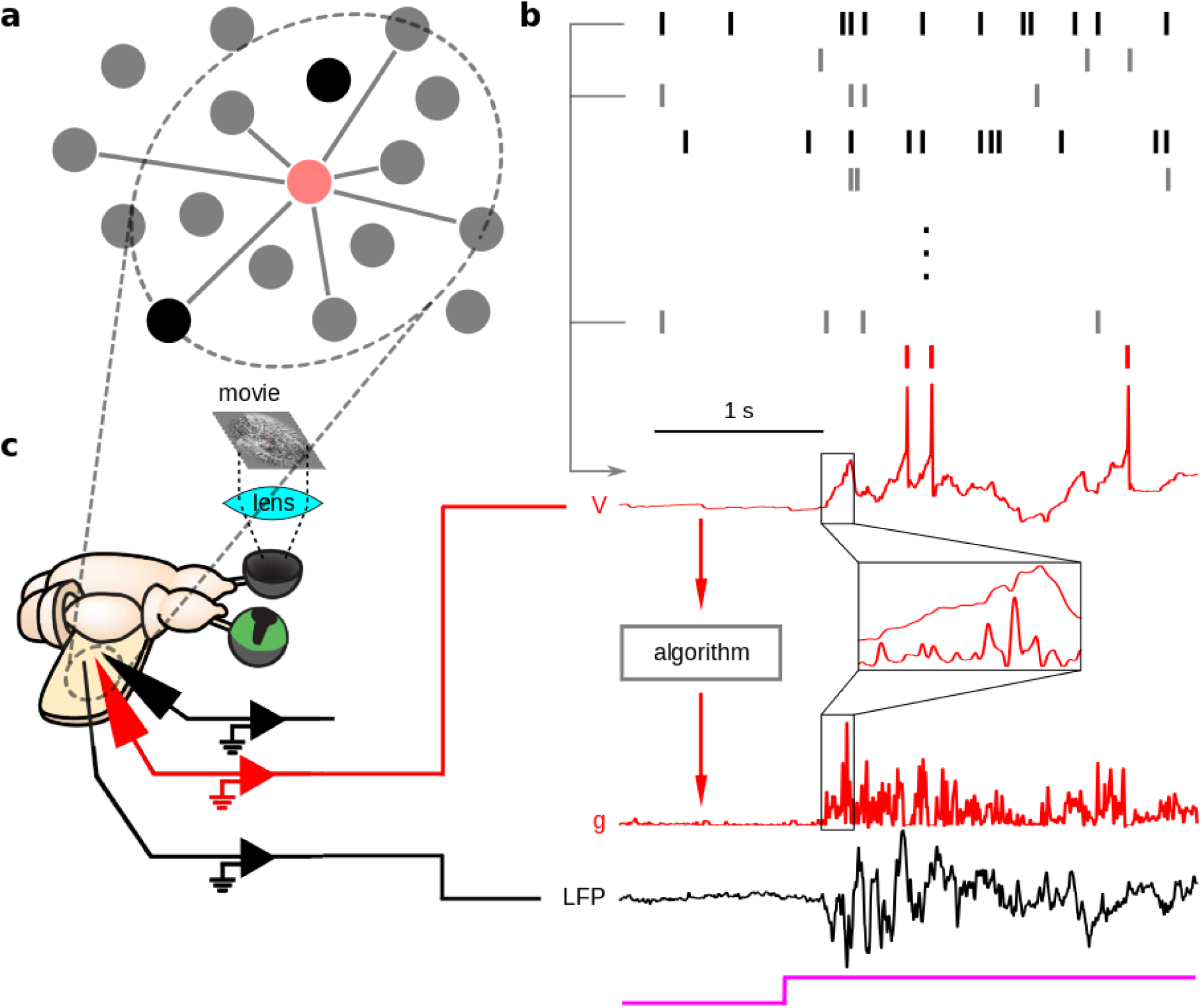
Individual neurons subsample the cortex, and provide a spike-rate-independent measure of cortical sensory responses. (a) Cortical neurons are primarily sparse-spiking units (low opacity circles), and each neuron subsamples the cortex by receiving synaptic inputs from a large, biologically relevant subpopulation. (b) While high-spike-rate neurons (high opacity rasters) alone provide reliable statistics for analysis of spiking responses, the subthreshold activity of a randomly-selected neuron (e.g., red voltage trace corresponding to red rasters) communicates information about the time course of presynaptic spiking activity. (c) Left: We recorded the subthreshold membrane potentials of cortical pyramidal neurons, as well as the nearby local field potential (LFP) while presenting movies to the retina in the turtle eye-attached wholebrain *ex vivo* preparation. Right: We used an algorithm (see Methods) to infer the excitatory synaptic conductance (g) from V, which gave a more detailed view of synaptic activity (inset). We investigated the nature of the variability in g, and its codupling with that of the simultaneously-recorded LFP.

We addressed the third question by implementing a small-world network of leaky integrate-and-fire neurons, subject to Poisson process external inputs and synaptic depression with recovery. This model reproduces three experimentally-observed aspects of evoked activity: large across-trial response variability, diverse g-LFP coupling across populations of nearby neurons, and the evoked coupling dynamic. These response properties are largely determined by the distribution of synaptic weights and the network state, which is highly sensitive to such network parameters as spatial clustering, synaptic time constants, and adaptation.

Together, our results provide a clearer picture of the subthreshold coordination dynamics corresponding to suprathreshold response variability and population coupling in cortex. Moreover, they implicate specific anatomical and emergent network properties that shape cortical variability and coordination during sensory processing.

## Results

### Visual stimulation evokes significant increases in cortical activity

To quantify the response variability of synaptic inputs and its coupling with that of the local population, we recorded the membrane potential (V) from 39 pyramidal neurons in visual cortex of the turtle *ex vivo* eye-attached whole-brain preparation during visual stimulation of the retina (**Figure 1c**). For 21 of these neurons, we also recorded the nearby LFP, which has been shown to be a reliable estimator of local synaptic activity(Haider et al. 2016). We then used a recently-developed algorithm(Yaşar, Wright, and Wessel 2016) to estimate the excitatory synaptic conductance (g) from V (**Figure 1d**, and see Methods).

Ongoing activity in turtle visual cortex was relatively quiescent, typically with infrequent postsynaptic potentials at the level of the membrane potential, and little to no baseline LFP activity (**Figure 1d, 2a**). On a minority of trials, this quiescent activity was interrupted by spontaneous “bursts” of activity lasting up to hundreds of milliseconds that were qualitatively similar to visual responses (**Figure 2a, d**). Visual stimulation evoked barrages of postsynaptic potentials, and large fluctuations in the nearby LFP (**Figure 1d, 2a**), with orders-of-magnitude increases in average power for both g and LFP (population-averaged relative power <rP>=3632.7 ± 3538.0, mean ± s.e.m., **Figure 2b, c**, <rP_LFP_> = 1902.9 ± 1350.7, data not shown, transient). Response amplitudes (**Figure 2a**) and power (**Figure 2b, c**) decreased from transient to steady-state, despite persistent visual stimulation (<rP> = 1251.9 ± 962.5, steady-state; P = 6.06×10^-8^ for transient – steady-state comparison, Wilcoxon signed-rank test; <rP_LFP_> = 557.9 ± 449.1, steady-state; P = 1.2 × 10^-3^ for transient – steady-state comparison, Wilcoxon signed-rank test).

**Figure 2.**
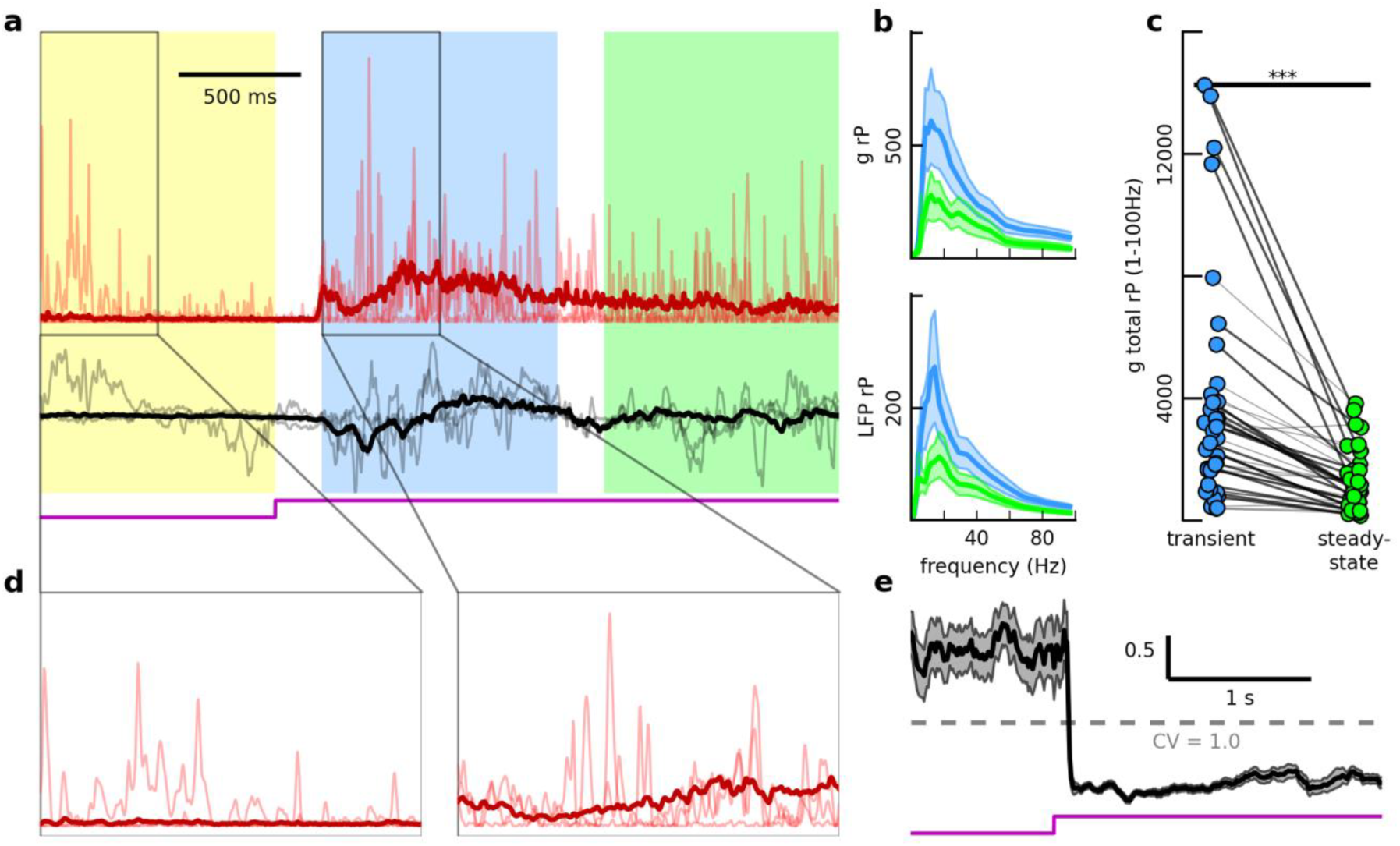
Visual stimulation evokes increases in synaptic activity, and responses are highly variable across trials. (a) Inferred excitatory synaptic conductance (g, red) and measured LFP (black) for three trials (low opacity), and average across 32 trials (high opacity). Colors indicate ongoing (yellow), transient (blue), and steady-state (green) epochs (see Results and Methods). (b) Relative power spectra (mean ± bootstrap intervals, see Methods) for g (top) and LFP (bottom) for transient (blue) and steady-state (green) epochs, for example pair in (a). (c) Total relative power (1 – 100 Hz) for 39 cells. Each blue (green) dot represents the across-trial mean relative power for one cell during the transient (steady-state) epoch. High opacity lines connecting dots indicate significant change across epochs (P < 0.05, bootstrap comparison test, see Methods). Asterisks above line connecting epochs indicate P < 1 × 10^-3^ (Wilcoxon signed-rank test). (d) Close-up view of ongoing (left) and evoked (right) synaptic activity. (e) Coefficient of variation (CV) as a function of time for 39 cells (mean ± s.e.m.). Dashed line indicates CV = 1.0.

### Visual responses are highly variable across trials

For a given cell and nearby LFP, the across-trial average responses to a given stimulus displayed clear temporal structure (**Figure 2a**). Still, responses were highly variable across stimulus presentations; single-trial fluctuations were large relative to the mean response, with the across-trial variability increasing along with the across-trial average activity (**Figure 2a, d**). To determine the relationship between the variability and the average response, we calculated the scaled variability, or coefficient of variation (CV), as a function of time, for the population of all cells (see Methods). While variability of evoked activity was larger than that of ongoing activity (**Figure 2d**), the population-averaged CV (<CV>) decreased after stimulus onset, and slowly recovered (**Figure 2e**). Using the windows of activity defined above, we found that this initial decrease was significant (<CV> = 1.83 ± 0.13 ongoing, 0.22 ± 0.04 transient, P = 1.74 × 10^-16^ for ongoing – transient comparison, Wilcoxon signed-rank test), and that <CV> increased significantly from transient to steady-state (<CV> =, 0.36 ± 0.06 steady-state, P = 1.74 × 10^-16^ for transient – steady-state comparison, Wilcoxon signed-rank test), but remained significantly smaller than during ongoing activity (P = 1.74 × 10^-16^ for ongoing – steady-state comparison, Wilcoxon signed-rank test).

### Additive and multiplicative noise contribute to response variability

We next investigated the nature of this single-trial variability. Even when a single-trial response deviates significantly from the mean response, it may follow a very similar (or in the extreme case, an identical) time course. This would indicate a high degree of “multiplicative noise”: a uniform modulation of the presynaptic population’s sensory response. Alternatively, in the case of purely “additive noise”, the single-trial time series fluctuates randomly about the average, consistent with noise amplitudes that vary across members of the presynaptic population.

To address this question, we first binned each single-trial inferred conductance (summing over 100 ms bins, resulting in 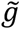), and then calculated the across-trial average binned conductance 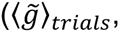 **Figure 3a**, and see Methods). By visual inspection, it was evident that individual responses contained both additive and multiplicative noise (**Figure 3a**). For example, a typical response that was somewhat “enveloped” by the average time course (indicating multiplicative noise) also tended to possess small, random fluctuations about the mean, or in some instances larger deviations away from the mean (**Figure 3a**, trial 3, steady-state epoch), examples of additive noise. To quantify the contributions of each component, we regressed 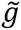 onto 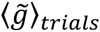 for each trial, and took the across-trial median R^2^ value for each cell and epoch. For a given cell, the across-trial average was a relatively poor predictor of the single-trial response (see example cell in **Fig 3b**); across the population, the average response explained only 28.1 ± 13.9% of the variance in individual trials for the transient epoch (across-cell average explained variance <R^2^> = 0.28 ± 0.14, **Figure 3c**). The explained variance was even lower during the steady-state (<R^2^> = 0.17 ± 0.15, **Figure 3c**), decreasing significantly from that of the transient epoch (P = 1.5 × 10^-3^ for transient – steady-state comparison, Wilcoxon signed-rank test). Evidently, single-trial responses contained substantial amounts of both multiplicative and additive noise, with additive noise dominating. In addition, the reliability of a single-trial response (as measured by its relationship to the across-trial average) diminished over the duration of the response.

**Figure 3.**
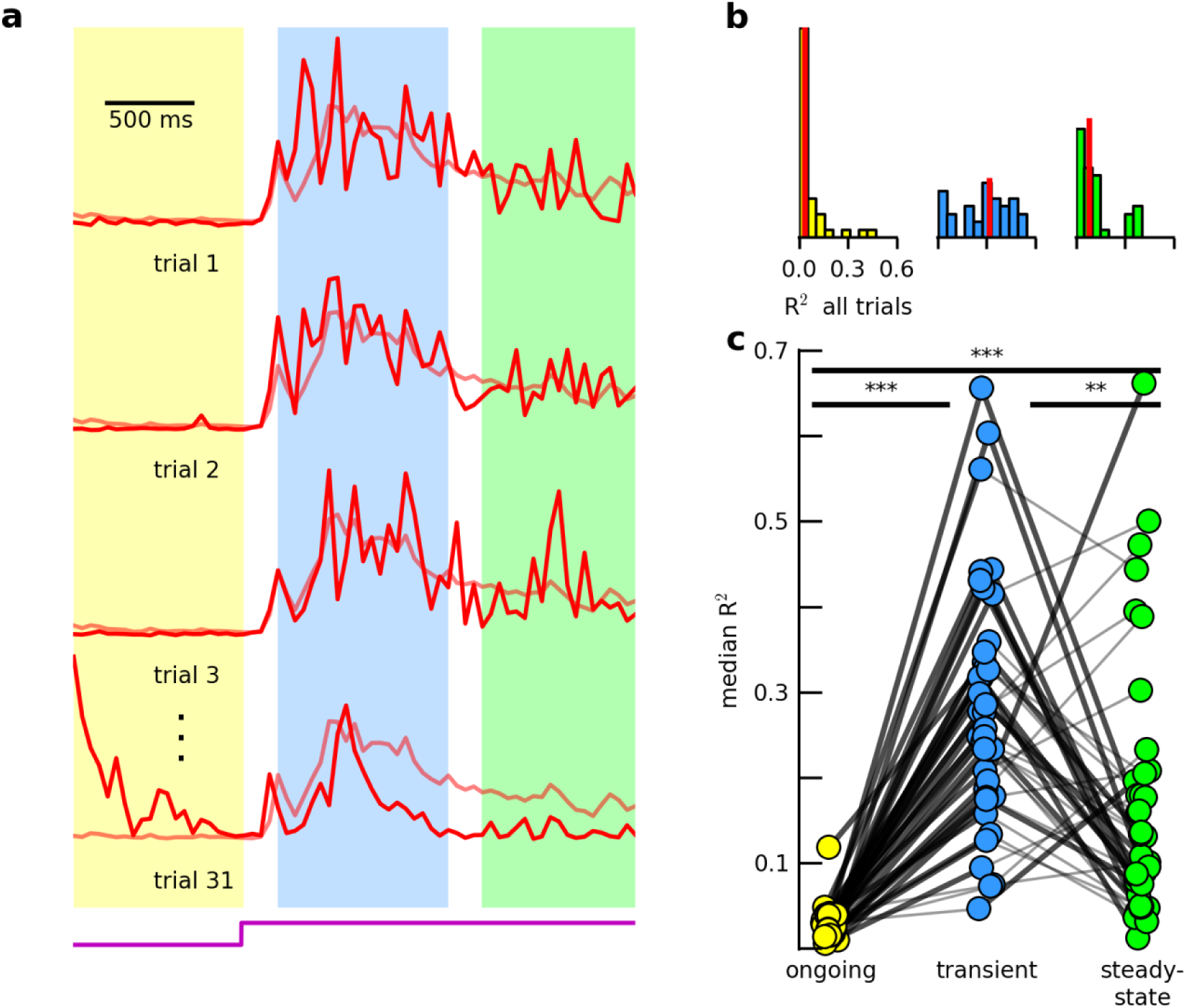
Single-trial variability is a mix of additive and multiplicative noise. (a) Inferred excitatory synaptic conductance, integrated over a 50 ms sliding window (with no overlap) for individual trials (high opacity), and across-trial average (low opacity, see Methods). (b) Across-trial distribution of R^2^ values resulting from linear regression of single-trial response onto average response for ongoing (left), transient (center), and steady-state (right) epochs. Red vertical lines indicate medians. (c) Across-trial median R^2^ values for 39 cells, for each epoch. Dot colors, connecting line opacities, and asterisks as in 2(b), with ** indicating 0.001 < *P* < 0.01, Wilcoxon signed-rank test.

### Correlated variability amplitude transiently increases following visual stimulation

Single-neuron response variability of this magnitude has the potential to profoundly influence sensory coding, provided it is significantly coupled across a population of neurons(Abbott and Dayan 1999; Averbeck, Latham, and Pouget 2006; Shadlen and Newsome 1998). We quantified this “population coupling” for 21 cells by calculating the single-trial residual responses for the estimated conductance (*gr*, the single-trial time series with the across-trial average time series subtracted) and the nearby LFP (**Figure 4a**) and calculating the Pearson correlation coefficient for residual pairs for each trial and epoch (see Methods).

**Figure 4.**
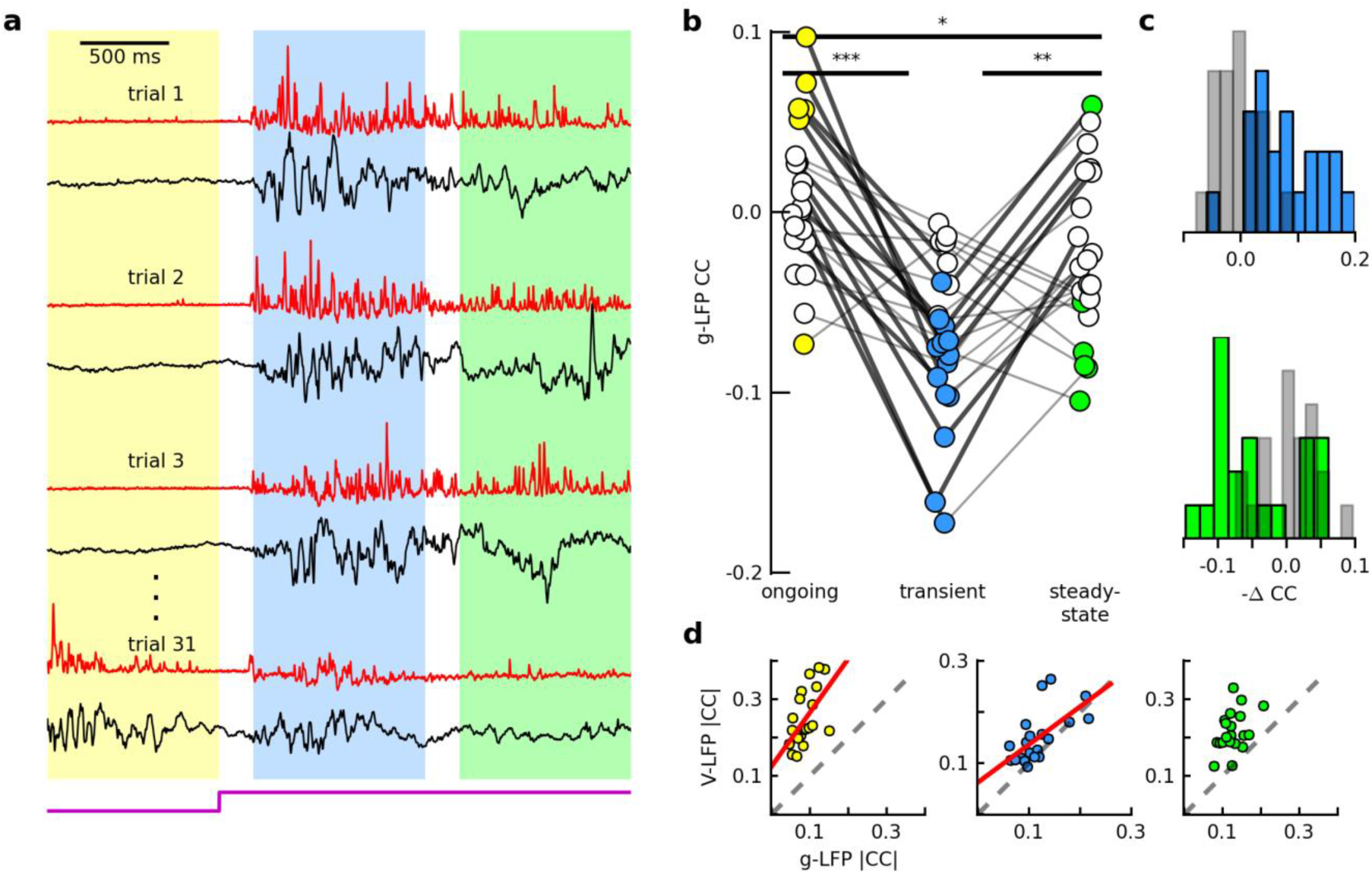
Synaptic input correlated variability transiently increases with visual stimulation. (a) Residual traces for g (red) and LFP (black) for multiple trials. (b) Across-trial average Pearson correlation coefficient for g and LFP residual traces, for 21 g-LFP pairs. Each dot indicates the across-trial average CC value for a given g-LFP pair. Filled dots indicate significant average values (P < 0.05, bootstrap comparison to shuffled data, see Methods). Connecting lines and asterisks as in 3(c), with * indicating ≤ 0.01<0.05, Wilcoxon signed-rank test. (c) Top: Distribution of change in across-trial average CC values (multipled by ‐1) from ongoing to transient, for 21 g-LFP pairs (with results for shuffled data shown in gray). Bottom: same, but for transient to steady-state. (d) Absolute value of across-trial average CC for V and LFP vs. that for g and LFP for 21 cell-LFP pairs, for the ongoing (left), transient (center), and steady-state (right) epochs. Dashed line is unity line, and red line indicates significant linear regression fit. V-LFP |CC| values were significantly larger than g-LFP |CC| values for all epochs (P = 8.86 × 10^−5^ ongoing, P = 2.2 × 10^−3^ transient, P = 8.9 × 10^-5^ steady-state, Wilcoxon signed-rank test). Values were significantly related for the ongoing (r^2^ = 0.31, P = 0.01 linear regression) and transient (r^2^ = 0.40, P = 0.0029) epochs, but not the steady-state (r^2^ = 0.18, P = 0.06).

For a given stimulus condition, the trial-averaged correlation coefficient (CC) was broadly distributed across the population (**Figure 4b**). During ongoing activity, CC was significantly nonzero for seven of 21 pairs (P < 0.05, comparison to shuffled data using Wilcoxon signed-rank test, see Methods). With visual stimulation, the population of pairs became more anti-correlated (**Figure 4b**); CC amplitudes increased significantly for 10 pairs (P < 0.05, across-epoch bootstrap comparison) and the population average decreased significantly (such that the amplitude increased; <CC> = 0.009 ± 0.04 ongoing, P = 0.50 for comparison to shuffled data; <CC> = −0.07 ± 0.04 transient, P = 1.1 × 10^−4^ for comparison to shuffled data; P = 1.9 × 10^−4^ for ongoing – transient comparison, Wilcoxon signed-rank test, **Figure 4b, 4c, top**). During this transient epoch, CC was significantly nonzero for 14 pairs (P < 0.05, comparison to shuffled data). This elevated level of coordination soon relaxed: from transient to steady-state, CC amplitudes significantly decreased for 5 pairs (P < 0.05, across-epoch bootstrap comparison), such that CC was significantly nonzero for 7 pairs (P < 0.05, comparison to shuffled data), and the population average increased significantly toward zero (<CC> = −0.02 ± 0.05 steady-state, P = 0.005 for transient – steady-state comparison, Wilcoxon signed-rank test) to values that were not significant across the population (P = 0.11 for comparison to shuffled data, **Figure 4b, 4c, bottom**).

These results suggest that the across-trial variability in evoked synaptic inputs to an individual neuron is, on average, coupled to that of other neurons in a nearby population in the early response phase. Moreover, the strength of this coupling is highly variable across cells. Coupling is not static, however; while response reliability decreases from the early to the late phase of the visual response (**Figure 3**), the coupling strength does as well (**Figure 4b, c**), suggesting that large single-trial fluctuations in the late response are more effectively “averaged out” across a large population.

### The dynamics of g-LFP CC are consistent with known excitation-inhibition dynamics

When combined with what is known of the recorded signals and excitatory-inhibitory dynamics, these g-LFP CC results provide a means for testing the validity of the inferred excitatory synaptic conductance. First, V and LFP are both a mix of excitation and inhibition (while our inferred conductance excludes inhibition), so for a given cell, V-LFP CC should be larger than g-LFP CC in a given window of activity. Second, because excitatory currents make a large contribution to V and LFP, and because inhibitory currents generally tend to track excitatory currents(Atallah and Scanziani 2009; Isaacson and Scanziani 2011; Wehr and Zador 2003), g-LFP CC and V-LFP CC should be related.

Our results largely satisfied these predictions. First, for each epoch of activity, across the population of cells, CC amplitudes were larger for V-LFP than for g-LFP (**Figure 4d**). This was also true when V and LFP were filtered in the gamma band (thus removing the shared, slow fluctuation to isolate the fast activity resulting from high-frequency synaptic inputs(Hasenstaub et al. 2005; Nowak, Sanchez-Vives, and McCormick 1997; Poulet and Petersen 2008), data not shown). Second, g-LFP and V-LFP CC amplitudes were significantly related for the ongoing and transient epochs for both 100 Hz low-pass (**Figure 4d**) and gamma band (data not shown) activity. There was also a positive relationship in the steady-state, but the trend was not significant (P = 0.059, **Figure 4d, right**). As such, these results provide further evidence for the reliability of the estimation algorithm.

### Network properties shape response variability and g-LFP correlated variability

We next sought to infer the relative contributions to the observed response properties from the stimulus and the thalamocortical network. Specifically, we asked which aspects of the experimentally-observed phenomena could be reproduced by a model network subject to random external inputs (mimicking the stimulus), and what network parameters were relevant to these results.

We implemented a model network similar to that described previously (N. C. Wright, M. Hoseini, R. Wessel, unpublished observations, **Figure 5a**). The network consisted of 800 excitatory and 200 inhibitory leaky integrate-and-fire neurons. Excitatory – to – excitatory connections had small-world connectivity (3%), and all other connections were random (3% excitatory – to – inhibitory, 20% inhibitory – to – excitatory and inhibitory – to – inhibitory). Nonzero synaptic weights were drawn from a beta distribution with mean value 1.0, which approximated the “constellation-like” connectivity in cortex(Cossell et al. 2015). All neurons received Poisson process external inputs, and the stimulus was modeled as an increase in external input rate. The external drive was unique across neurons and trials during the ongoing epoch. After stimulus onset, the external drive was a mix of two components: one that was unique across neurons, but identical across trials (with proportionality constant 0.75) and one that was unique across both neurons and trials (with proportionality constant 0.25). Because visual stimulation reliably evoked strong LFP oscillations (**Figure 2a**, and see Shew, et al. 2015), we selected a set of synaptic rise and decay times that were consistent with network spike rate oscillations in response to strong external drive (**Figure 5b**). Each synapse depressed and slowly recovered in response to a presynaptic spike. We modeled the LFP as the sum of all synaptic currents(Atallah and Scanziani 2009; Destexhe 1998) to a subpopulation of 100 neighboring excitatory neurons. We selected 40 neurons from the geometric center of this population for analysis of excitatory conductances (**Figure 5b**, and see Methods).

**Figure 5.**
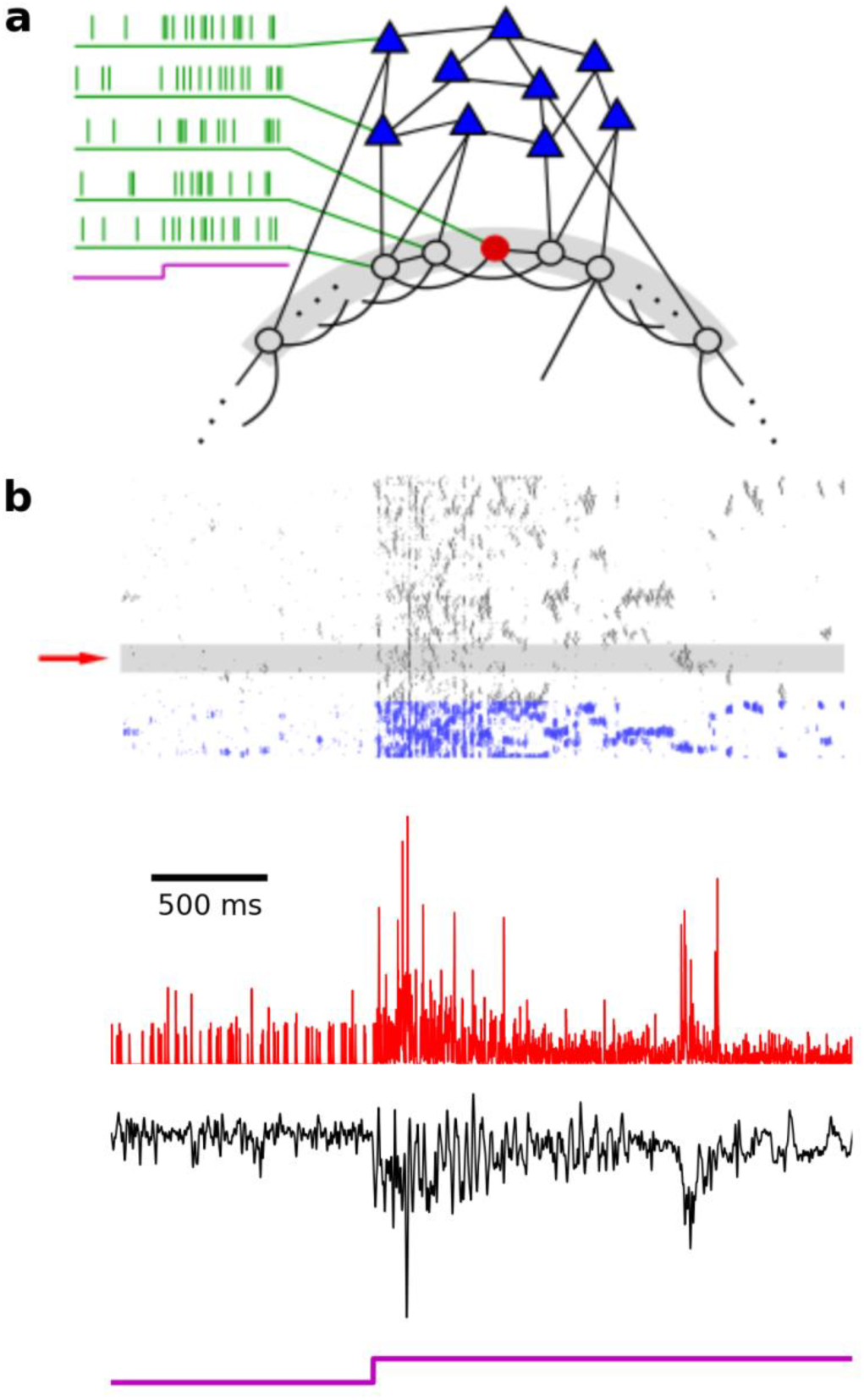
Model overview. (a) We implemented a model network of 800 excitatory and 200 inhibitory leaky integrate-and-fire neurons, all subject to Poisson process external inputs. Excitatory-excitatory connections had small-world connectivity, and all other connections were random. (b) Network parameters were tuned to give spike rate oscillations in the inhibitory (blue) and excitatory (black) populations in response to strong external drive. The LFP was modeled as the sum of synaptic currents to a subset of 100 neighboring neurons (gray region, and single trial in black below). The excitatory synaptic conductance was selected for neurons near the geometric center of this subset (single trial in red below, corresponding to neuron indicated by red arrow).

As in experiment, g and LFP varied considerably across trials (**Figure 6a**), despite the stimulus being primarily the same across trials (see Methods). As with our experimental data, we quantified the dynamics of this variability by calculating the scaled variability (CV) over time. The CV dynamics were determined by both the statistics of the external drive and by network properties. When external drive during the ongoing epoch was sufficiently strong to cause sparse network spiking, CV for the total excitatory synaptic conductance to network neurons hovered near 1.0 (<CV> = 0.95 ± 0.25, **Figure 6b**). This value greatly exceeded that of the external inputs alone (<CV> = 0.15 ± 0.09, **Figure 6b**), which was due to the highly variable distribution of nonzero synaptic weights (**Figure 6c, top**). With stimulus onset, CV for external inputs decreased by design (to <CV> = 0.004 ± 0.01), and CV for total excitatory conductance initially did as well (<CV> = 0.40 ± 0.16 for transient epoch, P = 4.27 × 10^-18^ for ongoing – transient comparison, Wilcoxon signed-rank test). This decrease in CV was due in part to the concerted increase in external drive, and in part to the stimulus possessing a component that was identical across trials (**Figure 6c, middle**). Over the course of hundreds of milliseconds, CV for total excitatory conductance recovered to nearly that of the ongoing epoch (<CV> = 0.81 ± 0.17 for steady-state epoch, P = 1.80 × 10^-16^ for transient – steady-state comparison, P = 1.1 × 10^−4^ for ongoing – steady-state comparison, Wilcoxon signed-rank test), which was an exaggeration of the experimental scaled variability dynamic observed here (**Figure 2e**) and elsewhere(Churchland et al. 2010). Synaptic depression mediated this recovery (**Figure 6c, bottom**). Thus, CV values and dynamics depended on the distribution of synaptic weights, the across-trial reliability of external inputs, and synaptic adaptation.

**Figure 6.**
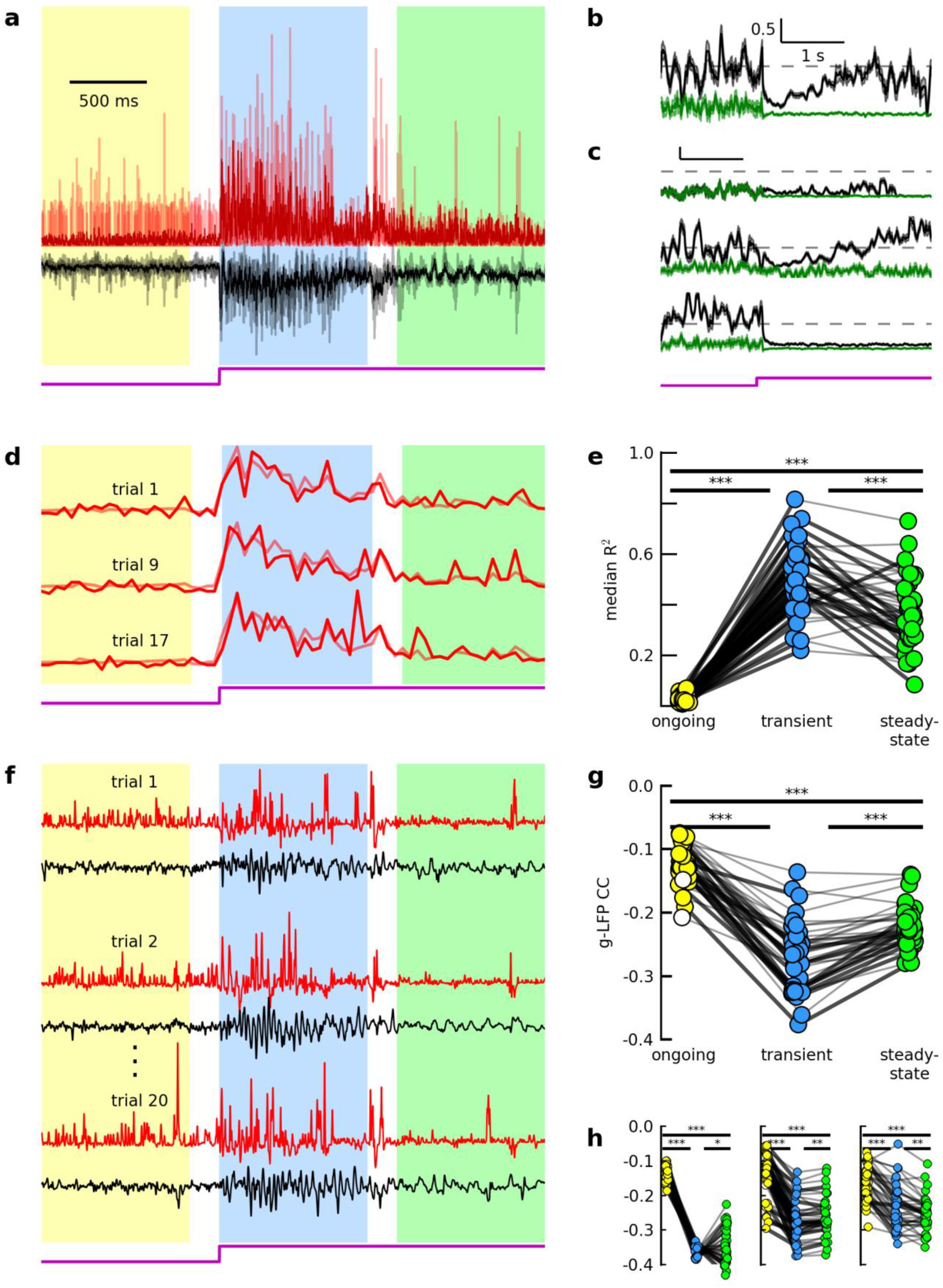
A model network qualitatively reproduces the experimental results. (a) excitatory synaptic conductance for one model neuron (g, red) and nearby LFP (black) for three trials (low opacity), and average across 20 trials (high opacity). Colors indicate ongoing (yellow), transient (blue), and steady-state (green) epochs. Example cell is located at the geometric center of the pool defining the LFP (see Results and Methods). (b-c) A qualitative reproduction of experimental CV(t) depended on the synaptic weight distribution, the nature of the stimulus, and synaptic depression. (b) Coefficient of variation (CV) as a function of time (mean ± s.e.m.) for 40 cells randomly-selected from the network, for total excitatory synaptic conductance (black), and for external excitatory conductance (green). Dashed line indicates CV = 1.0. (c) CV for alternate model versions. Top: network with binary synaptic weights. Middle: network subject to unique stimulus on each trial. Bottom: network without synaptic adaptation. Scale bar and dashed line same as in (b). (d) Excitatory synaptic conductance for one model neuron, integrated over a 50 ms sliding window (with no overlap) for individual trials (high opacity), and across-trial average (low opacity, see Methods). (e) Across-trial median R^2^ values for 40 cells randomly-selected from network, for each epoch. Dot colors, connecting line opacities, and asterisks as in 3(c). (f) Residual traces for g (red) for one model neuron and nearby LFP (black) for multiple trials. (g-h) A qualitative reproduction of the experimental g-LFP dynamic depended on the synaptic weight distribution, synaptic depression, and network oscillations. (g) Across-trial average Pearson correlation coefficient for g and LFP residual traces, for 40 g-LFP pairs (where 40 cells are selected from geometric center of pool defining LFP). Each dot indicates the across-trial average CC value for a given g-LFP pair. Filled dots indicate significant average values (P < 0.05, bootstrap comparison to shuffled data, see Methods). Connecting lines and asterisks as in 5(f). (h) Same as in (g), for alternate model versions. Left: network with binary synapses (i.e., synaptic weights either 1 or 0). Middle: network without synaptic depression. Right: Asynchronous network.

The model qualitatively reproduced the contribution of additive and multiplicative noise to the total response variability. As in experiment, single-trial variability was a mix of both noise types (**Figure 6d**, compare to **Figure 3a**), and the relative contribution from additive noise increased from transient to steady-state (<R^2^> = 0.50 ± 0.13 ongoing, <R^2^> = 0.37± 0.13 steady-state, P = 7.6 × 10^-6^ for transient – steady-state comparison, Wilcoxon signed-rank test, **Figure 6e**, compare to **Figure 3c**). This decrease in response reliability was not related to synaptic depression (data not shown), suggesting single trial “errors” compounded over the duration of the response. Notably, the percent of single-trial variance explained by the average response in either epoch was smaller than the 75% predicted by the stimulus. This surplus variability is therefore due to the only other source of randomness in the model: the state of the intracortical synapses at stimulus onset (due to the variable external drive and intracortical synaptic depression during the ongoing epoch, see Methods).

As in experiment, we calculated correlated variability (CC) for g-LFP pairs (**Figure 6f**). The synaptic weight distribution strongly influenced g-LFP CC distributions. For each epoch, CC was broadly distributed across the population (**Figure 6g**). While some variability is to be expected from such a sparsely-connected network, CC distributions were far less variable in a network with binary synapses (but the same average synaptic weight, **Figure 6h, left**).

The dynamics of g-LFP CC depended on a variety of network parameters. We recently used a similar network to demonstrate the effects of coordinated spiking on gamma band membrane potential correlated variability (N. C. Wright, M. Hoseini, R. Wessel, unpublished observations). Briefly, when the external drive triggers network spike rate oscillations, excitatory synaptic inputs to a given neuron become strongly correlated with both excitatory and inhibitory inputs to other neurons (with a small lag between excitation and inhibition). This leads to strong membrane potential oscillations that are correlated across neurons, an effect that is not observed in an asynchronous driven network. This coordination dynamic is also manifested as an increase in g-LFP correlated variability from the ongoing to transient epoch (<CC> = −0.12 ± 0.03 ongoing; <CC> = -0.27 ± 0.05 transient; P = 3.57 × 10^−8^ for ongoing – transient comparison, Wilcoxon signed-rank test **Figure 6f**). Synaptic depression with slow recovery (see Methods) diminished network activity levels, and crucially, abolished large-scale coordinated spiking (**Figure 5b**). This had the effect of drastically reducing g-LFP CC amplitudes from transient to steady-state (<CC> = −0.22 ± 0.03 steady-state; P = 1.1 × 10^−7^ for transient – steady-state comparison, Wilcoxon signed-rank test, **Figure 6g**), despite persistent network activity (**Figure 5b, 6f**). When either synaptic depression was removed (**Figure 6h, middle**) or the network was tuned to remain asynchronous (**Figure 6h, right**, see Methods), changes in <CC> were much smaller across epochs, and did not qualitatively match the experimental results. As such, these results implicate emergent network oscillations – and the corresponding relevant anatomical network properties (i.e., synaptic time constants and synaptic depression) – in the determination of g-LFP CC dynamics.

Taken together, the model investigation points to the cortical network as the primary determiner of the experimentally-observed response variability and population coupling dynamics of synaptic inputs. Specifically, the model identifies synaptic clustering, time constants, and depression as extremely relevant anatomical properties.

## Discussion

To obtain a spike-rate-independent measure of single-neuron variability, and to measure its coupling with local population activity, we simultaneously recorded the membrane potential from pyramidal neurons and the nearby LFP in the turtle visual cortex during ongoing and visually-evoked activity (**Figure 1**). We estimated the excitatory synaptic conductance (g) from the membrane potential, and quantified the across-trial variability in g and correlated variability with the LFP. To identify relevant cortical network mechanisms, we implemented a small-world network of leaky integrate-and-fire neurons subject to external drive (**Figure 5**).

Studies spanning several decades have described the remarkable degree of variability in the sensory-evoked spiking responses of cortical neurons(Britten et al. 1993; Carandini 2004; Scholvinck et al. 2015). Certain aspects of this variability suggest it is shaped by the cortex itself. First, cortical variability surpasses that of the inputs from LGN(Scholvinck et al. 2015). Second, evoked variability, when scaled by the overall activity level, tends to be smaller than that of spontaneous activity across a variety of cortical areas and behavioral states, suggesting it is a property of large, recurrent networks(Churchland et al. 2010). Third, single-neuron spiking variability can be modeled as a mix of multiplicative and additive noise due to global cortical activity(Goris, Movshon, and Simoncelli 2014; Lin et al. 2015). Our experimental results agree with this “cortico-centric” view of response variability. We observed that individual neurons subsampling the cortex receive excitatory synaptic inputs that are extremely variable across stimulus presentations (**Figure 2a, d, Figure 3**), with scaled variability (CV) decreasing soon after stimulus onset (**Figure 2e**). The time course of the visual response revealed the presence of both additive and multiplicative noise in the spatiotemporal sequence of presynaptic firing (**Figure 3**). Finally, across a variety of stimuli, scaled variability (**Figure 2e**) and the contribution from additive noise (**Figure 3**) increased from transient to steady-state. That is, response reliability and the nature of the variability changed in a stimulus-independent manner. Each of these results was qualitatively reproduced by a model network subject to an extremely simple external drive (**Figure 5, 6**).

Partitioning synaptic input variability into additive and multiplicative noise essentially assigns a measure of influence to two types of network fluctuations: a uniform scaling of the entire presynaptic pool (i.e., multiplicative noise), and scaling that is independent across subpopulations within the presynaptic pool (i.e., additive noise). Our experimental results suggest both types of modulation are present in visual responses, with the latter dominating at the time scale considered (**Figure 3**). Moreover, the variability in the early response contains almost twice as much multiplicative noise as that in the late response (**Figure 3c**). While it is possible that this dynamic is due to hidden, temporally variable, extracortical influences, our simple model network gave a similar result (**Figure 6d, e**), and points to an alternative explanation: a sensitivity to conditions at stimulus onset, with small deviations leading to increasingly random fluctuations about the mean over the duration of the response. Such chaotic dynamics are a hallmark of balanced networks(Shadlen and Newsome 1998; Vreeswijk and Sompolinsky 1996). While at first glance this seems extremely disadvantageous to sensory coding, the balanced regime has other advantages, including fast responses to changes in external stimuli(Vreeswijk and Sompolinsky 1996), effective signal propagation(Vogels and Abbott 2005), and maximized information capacity(W. L. Shew et al. 2011).

Previous work has shown that population coupling based on spiking activity is broadly-distributed across cells, which may reflect the degree to which a given neuron samples the local population(Okun et al. 2015), and the structure of that connectivity(Pernice et al. 2011). In agreement with this, we found that g-LFP correlated variability was broadly-distributed across cells for a given stimulus condition (**Figure 4b**). Our model results further reinforce the hypothesis that connectivity underlies this variability: CC distributions were broadly-distributed for relatively realistic, “constellation-like” synaptic weight distributions (**Figure 6g**), but narrowly-distributed for binary synapses (**Figure 6h, left**). These distributions also shaped the dynamics of scaled variability (**Figure 6b, c, top**). Evidently, cortical connectivity patterns are manifested in the response variability and coordinated variability of synaptic activity, which likely reflect the response properties of population spiking observed elsewhere.

Despite this apparent dependence on anatomical connectivity, previous work has shown that g-LFP coupling can increase with visual stimulation(Haider et al. 2006). In agreement with this, we found that g-LFP correlated variability amplitudes significantly increased when we presented movies to the retina (**Figure 4b, 4c, top**). This effect was short-lived, however; g-LFP CC amplitudes decreased significantly after the early response phase (**Figure 4b, 4c, bottom**), despite persistent synaptic and local population activity (**Figure 2a-c, 4a**). Was this a case of external stimuli imposing a particular coordination dynamic on the cortical circuitry, or was the thalamocortical system itself capable of exhibiting multiple coordination “states”? Our model results support the latter hypothesis: sufficiently strong external drive that is uncorrelated across neurons can trigger intrinsic network oscillations, which are characterized by elevated coupling of synaptic inputs, and the elevated coupling is abolished along with the oscillation (**Figure 5, 6**). These results are consistent with the observation that spontaneous fluctuations in cortical state can influence g-LFP(Haider et al. 2016) and spike-spike(Okun et al. 2015; Scholvinck et al. 2015) population coupling. In addition, this decrease in g-LFP coupling is consistent with the observed decrease in multiplicative noise (**Figure 3**); single-trial fluctuations become increasingly independent across neuronal subpopulations, which should be reflected in the nearby LFP(Deweese and Zador 2004). Moreover, our results build on this previous work by identifying specific anatomical features of cortex (e.g., synaptic time constants and synaptic adaptation) capable of influencing population coupling dynamics via emergent network phenomena.

According to one view of population coding, the decrease in g-LFP coupling in the late response may benefit cortical function: while steady-state activity is less reliable than that in the early response (**Figure 2e, 3**), these later fluctuations are more private, and therefore tend to average out across a neural ensemble(Zohary 1994). Our model results suggest this does not simply reflect a decrease in overall activity level, but rather the abolition of large-scale spike rate oscillations by synaptic depression (**Figure 5b, 6**). This is consistent with the emerging view that adaptation (in cortex and elsewhere) serves as much more than a modulator of activity levels, but is in addition a “knob” for fine-tuning a variety of functionalities(Benucci, Saleem, and Carandini 2013; Gutnisky and Dragoi 2008; Ollerenshaw et al. 2014; Woodrow L Shew et al. 2015; Zheng, Wang, and Stanley 2015).

One major limitation of our experimental work is the lack of direct measurements of inhibitory synaptic conductances, which are a key component of single-neuron and network-wide response properties. Inhibition represents a significant proportion of the total synaptic input to a given neuron(Haider, Häusser, and Carandini 2012), tends to be correlated across pairs of neurons(Hasenstaub et al. 2005), and the relative timing of excitatory and inhibitory currents may determine precise spike timing(Haider and McCormick 2009; Hasenstaub et al. 2005; Nowak, Sanchez-Vives, and McCormick 1997) and feature selectivity(Wilent and Contreras 2005). Furthermore, the inhibitory population is known to play a vital role in such emergent network phenomena as spike rate oscillations(Brunel and Wang 2003), and the excitation-inhibition balance may represent a fundamental aspect of the cortical code(Denève and Machens 2016). Our experimental approach can be modified to investigate inhibition. For example, excitatory conductances can be pharmacologically blocked, and/or the resting membrane potential of a patched neuron can be shifted to the excitatory reversal potential. In this case, the algorithm would provide a temporally-precise view of the inhibitory conductances responsible for the observed (relatively slow and convoluted) membrane potential deflections. More importantly, this approach can be combined with multi-whole-cell recording to simultaneously infer excitatory conductances in one cell, and inhibitory in another, similar to studies of evoked activity in rat somatosensory cortex(Okun and Lampl 2008), and spontaneous activity in rat hippocampus(Atallah and Scanziani 2009) and mouse thalamocortical slice(Graupner and Reyes 2013). This would be particularly useful in areas such as visual cortex, where responses can be complex and highly variable (thus limiting the utility of recording excitation and inhibition from one cell on alternating trials).

Here, we have treated each recorded neuron as a network sub-sampler, viewing the synaptic inputs as a record of presynaptic spiking activity. Of course, each neuron is also a contributing member of the network. And although the relationship is complex(Carandini 2004), the nature of the synaptic inputs is likely extremely relevant to that of output spiking activity(Doiron et al. 2016; Litwin-Kumar et al. 2011; Lyamzin et al. 2015). We therefore hypothesize that the broad g-LFP coupling distribution observed here corresponds to the diverse spike-spike population coupling observed elsewhere. A carefully-designed experiment can directly test this hypothesis. For instance, the resting membrane potential of a recorded neuron can be systematically manipulated to support spiking in some trials (which can be compared to local or global spiking activity), and limit it in others (which would be used to calculate g-LFP coupling). It is this ability to study both suprathreshold and subthreshold activity that makes whole-cell recordings so valuable in our quest to understand coordinated network activity(Doiron et al. 2016).

Taken together, our results demonstrate the highly variable nature of visually-evoked spatiotemporal spike patterns in cortical microcircuits. Further, they suggest that several properties of this variability are largely determined intracortically, and identify specific, highly relevant cortical parameters. Importantly, these cortical properties together lead to an adapted network state that is in many ways ideal for sensory processing. As such, these results contribute to a clearer picture of the effects of anatomical and emergent network properties on single-neuron sensory responses and network-wide function.

## Methods

### Surgery

All procedures were approved by Washington University’s Institutional Animal Care and Use Committees and conform to the guidelines of the National Institutes of Health on the Care and Use of Laboratory Animals. Fourteen adult red-eared sliders (*Trachemys scripta elegans*, 150-1000 g) were used for this study. Turtles were anesthetized with Propofol (2mg Propofol/kg), then decapitated. Dissection proceeded as described previously(Crockett et al. 2015; Saha et al. 2011; W. L. W. L. Shew et al. 2015). In brief, immediately after decapitation, the brain was excised from the skull, with right eye intact, and bathed in cold extracellular saline (in mM, 85 NaCl, 2 KCl, 2 MgCl_2_*6H_2_O, 20 Dextrose, 3 CaCl_2_-2H_2_O, 45 NaHCO_3_). The dura was removed from the left cortex and right optic nerve, and the right eye hemisected to expose the retina.

The rostral tip of the olfactory bulb was removed, exposing the ventricle that spans the olfactory bulb and cortex. A cut was made along the midline from the rostral end of the remaining olfactory bulb to the caudal end of the cortex. The preparation was then transferred to a perfusing chamber (Warner RC-27LD recording chamber mounted to PM-7D platform), and placed directly on a glass coverslip surrounded by Sylgard. A final cut was made to the cortex (orthogonal to the previous and stopping short of the border between medial and lateral cortex) allowing the cortex to be pinned flat, with ventricular surface exposed. Multiple perfusion lines delivered extracellular saline, adjusted to pH 7.4 at room temperature, to the brain and retina in the recording chamber.

### Intracellular Recordings

We performed whole-cell current clamp recordings from 39 cells in 14 preparations. Patch pipettes (4-8 MΩ) were pulled from borosilicate glass and filled with a standard electrode solution (in mM; 124 KMeSO_4_, 2.3 CaCl_2_-2H_2_O, 1.2 MgCl_2_, 10 HEPES, 5 EGTA) adjusted to pH 7.4 at room temperature. Cells were targeted for patching using a dual interference contrast microscope (Olympus). All cells were located within 300 microns of an extracellular recording electrode. Intracellular activity was collected using an Axoclamp 900A amplifier, digitized by a data acquisition panel (National Instruments PCIe-6321), and recorded using a custom Labview program (National Instruments), sampling at 10 kHz. The visual cortex was targeted as described previously(W. L. W. L. Shew et al. 2015)].

### Extracellular Recordings

We performed extracellular recordings at 12 recording sites in seven preparations. We used tungsten microelectrodes (MicroProbes heat treated tapered tip), with approximately 0.5 MO impedance. Electrodes were slowly advanced through tissue under visual guidance using a manipulator (Narishige), while monitoring for spiking activity using custom acquisition software (National Instruments). Extracellular activity was collected using an A-M Systems Model 1800 amplifier, band-pass filtered between 1 Hz and 20,000 Hz, digitized (NI PCIe-6231), and recorded using custom software (National Instruments), sampling at 10 kHz.

### Visual Stimulation

The visual stimulation protocol has been described previously (N. C. Wright, M. Hoseini, R. Wessel, unpublished observations). Briefly, visual stimuli were presented using a projector (Aaxa Technologies, P4X Pico Projector), combined with a system of lenses (Edmund Optics) to project images generated by a custom software package directly onto the retina. The stimulus was either a sustained gray screen, a naturalistic movie (“catcam”), a motion-enhanced movie (courtesy Jack Gallant), or a phase-shuffled version of the same movie (courtesy Jack Gallant and Woodrow Shew). In all cases, the stimulus was triggered using a custom Labview program (National Instruments).

For each cell and extracellular recording site, we selected one of the four stimuli listed above to present across all trials. The preparation was in complete darkness before and after each stimulus presentation. Stimuli lasted either 10 s or 20 s, and were shown at least 12 times, with at least 30 s between the end of one presentation and the beginning of the next.

### Processing of intracellular and extracellular voltage recordings

Raw data traces were down-sampled to 1000 Hz. We used an algorithm to detect spikes in the membrane potential, and the values in a 20 ms window centered on the maximum of each spike were replaced via interpolation. Finally, we applied a 100 Hz lowpass Butterworth filter. For extracellular recordings, we used a sine-wave removal algorithm to minimize 60 Hz line noise.

### Data included in analysis

For each extracellular recording site, we used visual inspection to determine the quality of the recordings. In general, we excluded recording sites from consideration if voltage traces displayed excessive 60 Hz line noise, low-frequency noise (likely reflecting a damaged electrode), or on average small response amplitudes relative to baseline.

For intracellular recordings, we also excluded some trials and cells. To include a given trial, we required the membrane potential to remain at or above the calculated inhibitory reversal potential from the beginning of the ongoing epoch to the end of the steady-state epoch. The inhibitory reversal potential was calculated using the Chloride concentrations in the intracellular and extracellular solutions, but because of partial transfer of intracellular solution to the cell interior, it was possible for the recorded membrane potential to drop below this value. This causes the conductance estimation algorithm to return a singularity. Rather than reset the inhibitory reversal potential to the minimum membrane potential value for such a trial, we took the more conservative approach of excluding the trial from consideration. We also excluded trials with excessive low-frequency artifacts or membrane potential drift. Finally, we considered only cells with twelve or more retained trials for analysis.

In some cases, an extracellular electrode remained at a single recording site while we performed whole-cell recordings either simultaneously or sequentially from multiple nearby cells. To calculate CC for a given g-LFP pair, we included only the trials in which both the intracellular and extracellular voltage were recorded and retained.

### Inferred excitatory conductance

The algorithm for obtaining an estimated excitatory synaptic conductance (g) from V for single trials has been described previously(Yaşar, Wright, and Wessel 2016). Briefly, our algorithm approximates a solution to the underdetermined equation

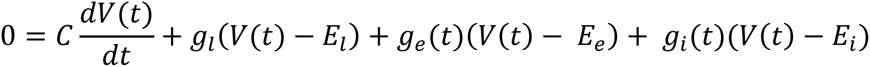

where *C* is the known membrane capacitance, *V*(*t*) is the measured membrane potential as a function of time, *E_e_* (*E_i_*) is the known excitatory (inhibitory) reversal potential, *E_l_* is the known leak reversal potential, *g_l_* is the known leak conductance, and *g_e_(*t*) (*g_i_*(*t*))* is the unknown excitatory (inhibitory) synaptic conductance. To estimate *g_e_*(*t*), we first introduce a mathematical construct *η*(*t*), which is defined according to

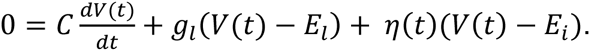

For each recording, we solve this equation for *η*(*t*). This attributes all membrane potential fluctuations to a single (unrealistic) inhibitory conductance. As such, *η*(*t*) contains negative values and rapid downward fluctuations that are due to the influence of excitatory currents on the membrane potential. Because conductance cannot have negative values, we then set the negative values in *η*(*t*) equal to zero, resulting in *ñ*(*t*) (previously called “non-negative *β*(*t*)” Next, we use linear interpolation to smooth out the rapid fluctuations in *ñ*(*t*). The output of this smoothing process is *ξ*(*t*), a smoother and therefore more realistic estimation of the inhibitory synaptic conductance. Finally, we substitute *ξ*(*t*) into the equation

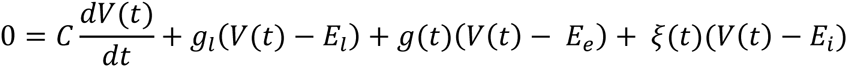

to obtain an estimation of the excitatory synaptic conductance (*g*). In general, this algorithm sacrifices knowledge about the inhibitory conductance to gain a better estimation of the excitatory conductance. Further, it capitalizes on the fact that excitatory currents are faster than – and therefore tend to interrupt – inhibitory currents.

We have made several improvements to the algorithm since introducing it. The original algorithm worked remarkably well on simulated membrane potentials. A recorded membrane potential, however, will contain high-frequency noise, which is removed by filtering (with, e.g., a 100 Hz Butterworth low-pass filter). This filtering process also leads to a smoother *ñ*(*t*). As mentioned above, detecting fast fluctuations in this signal is a critical step in the estimation process, and the algorithm’s performance was thus compromised by the filter (as evidenced by its application to filtered, noisy simulated membrane potentials). We therefore revised the criteria for detecting and replacing rapid fluctuations in *ñ*(*t*) (see Yasar, et al. 2016 for previous criteria). First, after calculating *ñ*(*t*), we obtained the time series *d*(*ñ*(*t*))/*dt.* We then determined each time *t’* at which *d*(*ñ*(*t*))/*dt* crossed a threshold of one negative standard deviation. This threshold optimized the algorithm’s performance when applied to noisy simulated data. Finally, we linearly connected the local maxima of *ñ*(*t*) immediately prior and posterior to *t’*.

When applying the algorithm to a membrane potential recording, the experimenter must estimate the resting membrane potential for that trial. An unrealistic choice will lead to spurious waveforms in the estimated conductance. We estimated the resting membrane potential for each trial by calculating the median membrane potential value during the quiescent activity in that trial. To isolate this quiescent activity, we first removed a window of activity starting at stimulus onset, and ending 6 s after stimulus offset. This resulted in either a 14 s or 24 s trace of “spontaneous” activity that was on average quiescent relative to that in the removed window. We then used an algorithm to detect spontaneous “bursts” of activity lasting at least 1 s in duration within the remaining trace, and removed these bursts. Finally, we took the median value (which is more robust to outliers than the mean) of the resulting trace to be the resting membrane potential for the corresponding visual stimulation trial.

### Coefficient of variation

The coefficient of variation (CV) is a scaled measure of variability: the standard deviation divided by the mean. For the set of all cells (N = 39), we calculated CV as a function of time (CV(t), **Fig 2e**) for the inferred excitatory conductance. First, we applied a 100 ms “box filter” to each g trace: for each time step, we replaced the value of the trace with the average value in a 100 ms window starting at that time step. We then advanced the window ten milliseconds, and repeated the process for the full length of the trace. Then, for each cell, we calculated the across-trial standard deviation and mean of the filtered traces as a function of time. This was done for the entire population, resulting in 39 (mean, standard deviation) ordered pairs for each time step. For each time step, we fit the set of means to the set of standard deviations using linear regression. The slope (standard error) of this fit was the coefficient of variation (s.e.m.) for the time step. To determine the significance of a change in CV across epochs, we compared the set of all CV values for one epoch with that from the other (e.g., the 100 values from the ongoing epoch and the 100 values from the transient) using the Wilcoxon signed-rank test.

### Correlated variability

For each single-trial time series X, the residual (Xr or deviation from the average activity) was found by subtracting the across-trial average time series from the single-trial time series:
*x*_*r*_ = *x* − 〈*x*〉_*trials*_

Residuals were then separated into three epochs: the ongoing epoch (defined to be the one second prior to the onset of visual stimulation), the transient epoch (200 to 1200 ms after stimulus onset), and the steady-state epoch (1400 to 2400 ms after stimulus onset; **Figure 4a**). For each g-LFP pair, the Pearson correlation between residuals was then calculated for each epoch and trial. The results were averaged across all trials, resulting in the trial-averaged correlated variability (CC) for each pair and epoch:

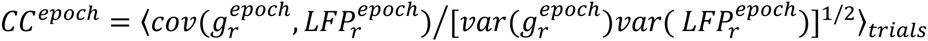

Significance tests for each pair and the population of pairs were applied as described below in “statistical analysis”.

### Power

For each trial and signal, we extracted a 4.4 s window of activity (with epoch windows and gaps between epochs as described above, plus 500 ms windows on each end to avoid filtering artifacts in the ongoing and steady-state epochs), and calculated the residual time series as described above. For each residual trace, we performed wavelet analysis in Matlab using software provided by C. Torrence and G. Compo(Torrence and Compo 1998) (available at URL: http://paos.colorado.edu/research/wavelets/). This resulted in a power time series for each cell, for multiple frequencies. For each frequency below 100 Hz, we averaged the time series across each epoch to obtain the average power at each frequency for each epoch. We then averaged across trials to obtain P^epoch^. For each cell, we also obtained the relative power spectrum (rP^epoch^) for the transient and steady-state epochs, defined to be the trial-averaged evoked spectrum divided by the trial-averaged ongoing spectrum (**Figure 3b**):

*_r_p^epoch^* = *p^epoch^ /p^ongoing^*

For each frequency, we calculated the bootstrap interval for the relative power as described below in “statistical analysis”.

### Network Models

To investigate the roles of network properties in our experimental results, we implemented a model network of 800 excitatory and 200 inhibitory leaky-integrate-and-fire neurons (**Figure 5a**). Excitatory-excitatory connections had small-world connectivity(Watts and Strogatz 1998) (with 3% connection probability), and all other connections were random (with 3% excitatory-inhibitory, and 20% inhibitory-excitatory and inhibitory-inhibitory connection probability). To generate each nonzero entry in the connection weight matrix 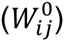 we drew a value from a beta distribution (over the interval [0.0, 1.0), with average value 0.1), and multiplied by 10.

The dynamics of the membrane potential (V) of each neuron evolved according

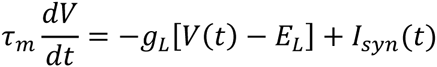

where the membrane time constant T_M_ = 50 ms (excitatory neurons), 25 ms (inhibitory), and the leak conductance g_L_ = 10 nS (excitatory), 5 (inhibitory). The leak reversal potential E_L_ for each neuron was a random value between -70 and -60 mV, drawn from a Gaussian distribution (to model the variability in resting membrane potentials observed across neurons in the experimental data). The reversal potentials for the synaptic current I_syn_(t) were E_GABA_ = -68 mV, and E_AMPA_ = 50 mV.

The synaptic current for each synapse type (between presynaptic neurons of type × and postsynaptic neurons of type Y) had three relevant time course parameters: delay (τ_LX_, that is, the lag between presynaptic spike time and beginning of conductance waveform), rise time (τ_RYX_), and decay time (T_DYX_). Synaptic conductances were modeled as products of time-varying gating variables (S_YX_) and maximum conductances (g_Yx_). Following a presynaptic spike at time 0, the gating variable dynamics were described by

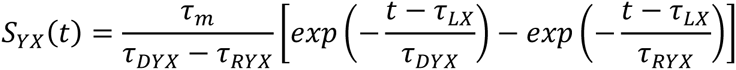

with time constants (in ms) τ_LE_ = 1.5, τ_REE_ = 0.2, τ_DEE_ = 1.0, τ_RIE_ = 0.2, τ_DIE_ = 1.0, τ_LI_ = 1.5, τ_RII_ = 1.5, τ_DII_ = 6.0, τ_REI_ = 1.5, τ_DEI_ = 6.0. Maximum conductance values (in nS) were gEE = 3.0, g_IE_ = 6.0, g_EI_ = 30.0, g_II_ = 30.0. In response to a presynaptic spike in neuron *j* at time 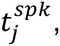, the weight (*W*_*ij*_) of a synapse connecting neurons *j* and *i* depressed and recovered according to

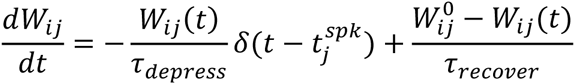

with depression time constant τ_depress_ = 300 ms and recovery time constant τ_recover_ = 2500 ms. Intracortical synapses were subject to depression for the entire simulation.

The spike threshold for each neuron was -40 mV. A neuron reset to -59 mV after spiking, and was refractory for 10 ms (excitatory) and 5 ms (inhibitory).

All excitatory and inhibitory neurons received Poisson external inputs. During “ongoing” activity, the external input rate to each neuron was 65 Hz, which was sufficiently high to cause intracortical spiking (**Figure 5b**). The ongoing external input was unique across cells and trials. The stimulus was modeled as a gradual increase to 375 Hz; the input rate was increased by 77.5 Hz at stimulus onset, and by an additional 77.5 Hz every 50 ms for 200 ms. This gradual increase provided more realistic excitatory conductances than did a single step function stimulus, but did not qualitatively impact the results. The post-stimulus external drive was composed of two components: one that was unique across cells and trials, and one that was unique across cells, but identical across trials, multiplied by proportionality constants 0.25 and 0.75, respectively. Thus, for the post-stimulus external drive to each neuron, 25% of the variance was explained by an input that was unique to each trial, and 75% was explained by an input that was identical across trials. The gating variables for external inputs had the same parameters as for excitatory-excitatory connections, and maximum conductances were gE = 6.0 nS. There was no thalamocortical synaptic depression during the ongoing epoch; the external drive during this window was simply used to generate stimulus-independent intracortical spiking, and thus treated as the “hidden” source triggering intrinsic events, as observed in experiment (**Figure 2a, d**).

Each trial was 4.4 s in duration, with stimulus onset at 1.7 s, and the time step was 0.05 ms. The ongoing epoch was defined to be 1200 ms to 200 ms before stimulus onset, the transient epoch 0 ms to 1000 ms after stimulus onset, and the steady-state epoch 1200 ms to 2400 ms after stimulus onset. The additional 500 ms at the beginning and end of each trial ensured there were no wavelet filtering artifacts in the ongoing and steady-state epochs.

We modeled the LFP as the sum of all synaptic currents(Atallah and Scanziani 2009; Destexhe 1998) to 100 neighboring neurons (**Figure 5b, d**), multiplied by a factor of -1 (to mimic the change in polarity between voltages measured intracellularly and extracellularly). The contribution of each neuron to the LFP was not distance-dependent. We then randomly selected 40 neurons from this subpopulation of 100 neurons, and used the excitatory synaptic conductances (**Figure 5c**) to generate 40 g-LFP pairs for g-LFP CC analysis (**Figure 6f-h**). For CV and R^2^ analysis (**Figure 6b-e**), we used 40 neurons randomly selected from the full population of 800 excitatory neurons.

### Statistical analysis

All statistical tests were performed using Python 2.7.

Before applying any significance test that assumed normality, we performed an omnibus test for normality on the associated dataset(s). This test compares the skew and kurtosis of the population from which the dataset was drawn to that of a normal distribution, returning a p-value for a two-sided chi-squared test of the null hypothesis that the data is drawn from a normal distribution. This test is valid for sample sizes of 20 or larger, and was implemented using scipy.stats.mstats.normaltest (documentation and references available at http://docs.scipy.org/doc/scipy-0.14.0/reference/generated/scipy.stats.mstats.normaltest.html). We report these p-values as the result of a “two-sided omnibus chi-shared test for normality”.

When asking whether a parameter of interest changed significantly across epochs for a population (e.g., whether the population-averaged CC for 21 g-LFP pairs changed significantly from the ongoing to transient epoch, see **Fig 4b**), we applied the Wilcoxon signed-rank test, which returns a p-value for the two-sided test that the two related paired samples (representing, e.g., the 21 (CC^ongoing^, CC^transient^) paired values) are drawn from the same distribution. This test assumes normality, and was implemented using scipy.stats.wilcoxon (documentation and references available at http://docs.scipy.org/doc/scipy/reference/generated/scipy.stats.wilcoxon.html).

To test whether a trial-averaged parameter of interest for one cell or electrode (e.g., CC, averaged over 15 trials for one cell) changed significantly from one epoch to another, we used a bootstrap comparison test. For each epoch of interest, we calculated the +/− 97.5% confidence intervals for the average value by bootstrapping (that is, resampling with replacement). If the bootstrap intervals for the two epochs did not overlap, we reported that the two sets of values were significantly different (p < 0.05).

When calculating correlations between a pair of signals in which at least one is slowly-varying, it is possible for broad autocorrelations to introduce spurious cross-correlations. This should be dealt with by either removing the broad autocorrelations (e.g., by “pre-whitening” the signals), or by accounting for their contribution to the cross-correlation. To avoid changing the temporal structure of the visual responses, we chose the latter approach. First, for each epoch and g-LFP pair, we randomly shuffled the trial order for one of the channels. We then calculated CC_shuff_ and the bootstrap interval for this shuffled data. The CC value for each pair and epoch was determined to be significant (with p < 0.05) if the bootstrap intervals for CC and CC_shuff_ data did not overlap. We indicate a significant CC value with a filled dot in the CC trajectory (**Figure 4b, 6g, h**). Finally, for a given epoch, we compared the sets of CC and CC_shuff_ values for the population of g-LFP pairs using the Wilcoxon signed-rank test (as described above for across-epoch comparisons of CC). The population average for unshuffled data was determined to be significant for p < 0.05. We repeated this second test using bootstraps intervals rather than the signed-rank test, with similar results (data not shown).

